# Cardiac Troponin I Directly Binds and Inhibits Mitochondrial ATP Synthase: a Noncanonical Role in the Post-Ischemic Heart

**DOI:** 10.1101/2023.02.03.526715

**Authors:** Aly Elezaby, Amanda J Lin, Vijith Vijayan, Suman Pokhrel, Luiz RG Bechara, Nicolai P Ostberg, Bruno B Queliconi, Juliane C Campos, Julio CB Ferreira, Bereketeab Haileselassie, Daria Mochly-Rosen

## Abstract

Cardiac troponin I (cTnI) is a sarcomeric protein critical to myocyte contraction. Unexpectedly, we found that some cTnI localized to the mitochondrial matrix in the heart, inhibited mitochondrial functions when stably expressed in non-cardiac cells and increased opening of the mitochondrial permeability transition pore under oxidative stress. Direct, specific, and saturable binding of cTnI to ATP synthase was demonstrated *in vitro*, using immune-captured ATP synthase, and in cells using proximity ligation assay. cTnI binding doubled F_1_F_0_ ATPase activity, whereas skeletal troponin I and several human mutant cTnI variants associated with familial hypertrophic cardiomyopathy did not. A rationally-designed ten amino acid peptide, P888, inhibited cTnI binding to ATP synthase, inhibited cTnI-induced increase in ATPase activity *in vitro*, and reduced cardiac injury following transient ischemia *in vivo*. We therefore suggest that mitochondria-associated cTnI may inhibit cardiac ATP synthase under basal conditions; pharmacological agents that release this inactivating effect of cTnI and thus preventing ATP hydrolysis during cardiac ischemia may increase the reservoir of functional mitochondria to reduce cardiac injury.

**Significance Statement:** Cardiac troponin I (cTnI) is a key sarcomeric protein involved in the regulation of myocardial contractility. We found that some cTnI is present in the mitochondrial matrix where it binds to ATP synthase, disrupting mitochondrial function; inhibition of the cTnI-ATP synthase interaction with a selective peptide inhibitor reduces cardiac dysfunction following ischemia and reperfusion injury. Several pathogenic cTnI mutations associated with hypertrophic cardiomyopathy do not affect ATP synthase activity, suggesting a potential mechanism that contributes to the diverse pathologies associated with these mutations.

## Introduction

The human heart produces and consumes an average of 6 kilograms of adenosine triphosphate (ATP) per day, relying primarily on mitochondrial metabolism for its energy demands^1^; in the healthy heart, 60-70% of ATP is utilized for myocyte contraction^2,3^. Actin and tropomyosin filaments form the contractile elements and are regulated by three associated proteins: troponin T, troponin C, and troponin I, which together form the troponin complex. The troponin complex attaches to tropomyosin, thus physically blocking and preventing muscle contraction. Upon calcium binding to troponin C, actin binds to myosin to enable the contraction of the myocyte^4^. While cytoskeletal elements such as actin and microtubule filaments interact with mitochondria to regulate mitochondrial fusion-fission^5–7^, trafficking^8,9^, membrane attachment^10^, and cellular metabolism^11–14^, whether contractile proteins have a direct role in regulating mitochondrial function is not known.

Myocardial infarction, the leading cause of cardiovascular death^15^, causes impaired mitochondrial function, bioenergetics and redox balance in cardiomyocytes, which ultimately results in cell death, tissue remodeling and the development of heart failure^16–18^. Under basal conditions, the majority of cardiac troponin I (cTnI) is myofibril-bound, but ~2-4% remains unbound to the contractile element^19^. During cardiac ischemia-reperfusion (IR) injury, cTnI phosphorylation induces structural alterations in the protein and increases its proteolysis, affecting muscle contractility^4,20,21^ concomitantly with reduced mitochondrial function^22^. Ischemia also induces the release of troponin subunits from the contractile elements, especially cTnI, and the increase in cTnI levels in the blood is used as a biomarker for myocardial injury in patients^19,23^.

In addition to its established structural role, non-cytoskeletal cTnI can localize to the nucleus^24^ where it regulates transcription^25,26^ and induces inflammatory signaling^27,28^ in heart failure. Furthermore, human cTnI mutations are associated with aberrations in mitochondrial structure and function in the heart through unclear mechanisms^29–31^, suggesting a potential direct connection between cTnI and mitochondrial functions. Here we identified how cTnI directly regulates mitochondrial functions and a means to inhibit this non-canonical role of cTnI to reduce cardiac injury following ischemic insult.

## Results

### cTnI localizes to mitochondria and regulates their function

To determine if cTnI may directly regulate mitochondrial functions, we first examined whether cTnI localizes to mitochondria. Using mouse hearts, we found that some endogenous cTnI associates with the mitochondria-enriched protein fraction under basal conditions, whereas another contractile protein, troponin C, did not (Figure 1A). To visualize cTnI localization, we transfected H9c2 cardiac myoblasts with a cTnI construct possessing fluorescent tags (GFP on the N-terminus and mCherry on the C-terminus). C-terminal mCherry fluorescence greatly overlapped with mitochondria labeled with MitoTracker (Figure 1B), especially in the cell periphery. Notably, the localization of N-terminal-labeled cTnI was cytosolic and punctated and not co-localized with the mitochondria, which may indicate that a portion of the N-terminal extension of cTnI is cleaved and doesn’t enter mitochondria, or that the GFP tag interferes with mitochondrial localization.

**Figure 1.**
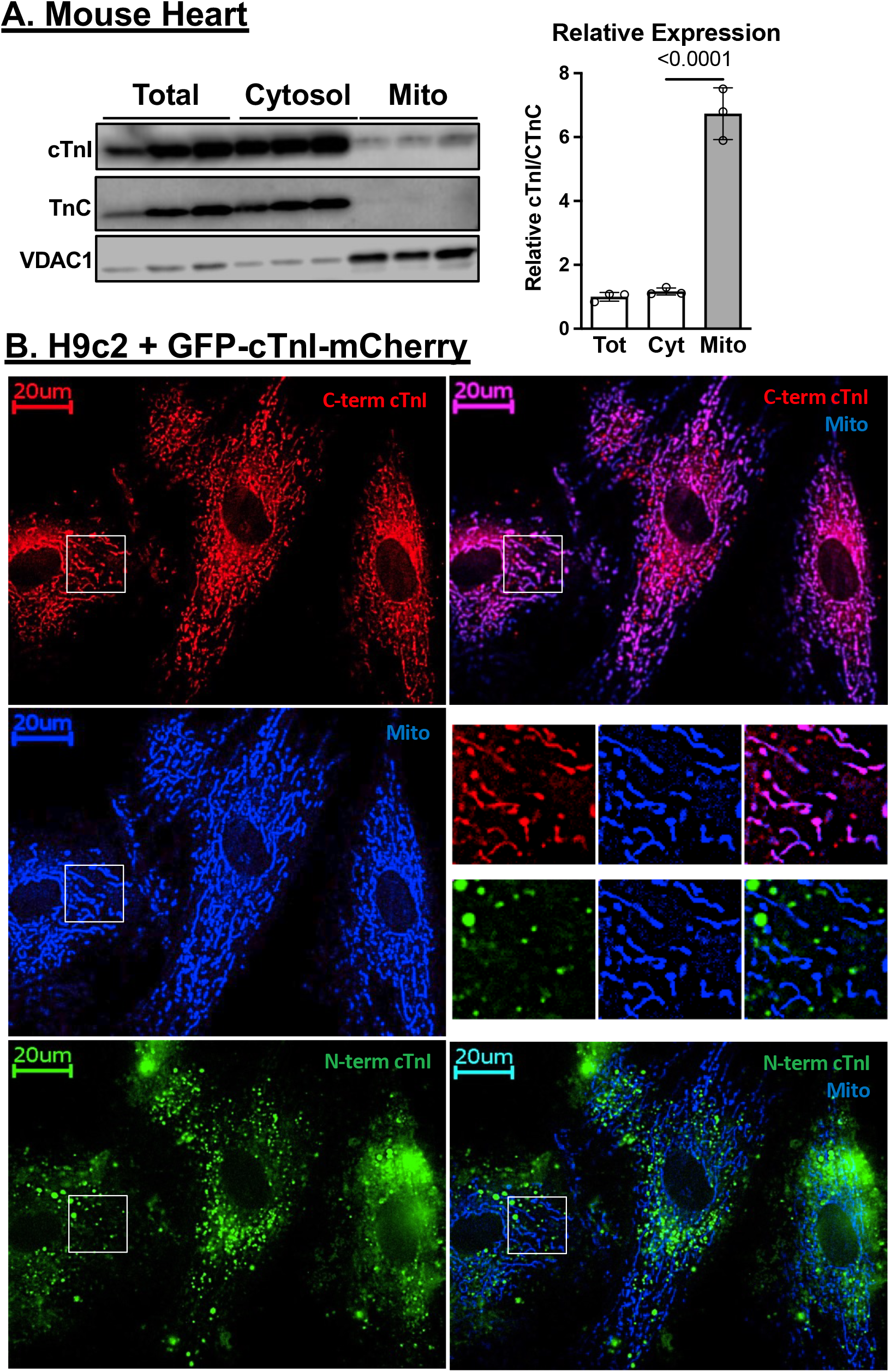
cTnI localizes to mitochondria. (A) *Left:* Total, mitochondrial and cytosolic protein fractions were isolated from mouse hearts, and western blot was used to determine the presence of cTnI. VDAC1 was used as mitochondrial protein control and troponin C (TnC) was used as a cytosolic protein control; n=3. *Right:* Quantification of immunoblot densitometry for cTnI normalized to TnC. (B) H9c2 rat cardiac myoblasts were transfected with a cTnI construct possessing N-terminal GFP and C-terminal mCherry tags. Images were taken for mCherry (red) on the C-terminus (*top left*), GFP on the N-terminus (*bottom left*), and mitochondria stained with Mitotracker (pseudo-color blue, *middle left*). Co-localization of C-terminal cTnI with mitochondria is observed (*top right*), but no co-localization of N-terminal cTnI is seen (*bottom right*).

We reasoned that if cTnI can localize to mitochondria independently of a contractile apparatus, this should also occur when cTnI is expressed in non-cardiac cells. We therefore engineered human embryonic kidney cells (HEK293T) to stably express full length cTnI (HEK-cTnI; Figure 2A, top). cTnI localized to the mitochondria-enriched protein fraction in HEK-cTnI cells (Figure 2A, bottom). Relative to control HEK cells, HEK-cTnI cells showed a 30% decrease in ATP level without a change in mitochondrial mass (Figure 2B). Using both FACS and fluorescence microscopy, we found that the mitochondrial membrane potential was lower in HEK-cTnI cells compared to control cells (Figure 2C). HEK-cTnI cells exhibited a decrease in basal (state II) and maximal (FCCP-uncoupled) oxygen consumption rates as well as in proton leak (state IV) (Figure 2D), suggestive of an overall decrease in mitochondrial respiration. Finally, when exposed to an oxidant stressor, H_2_O_2_, HEK-cTnI cells had a greater increase in the rate of opening of the mitochondrial permeability transition pore (mPTP) as measured by the rate of decrease in TMRM fluorescence in the presence and absence of the mPTP inhibitor cyclosporine A (CsA) (Figure 2E). These data suggest that cTnI expression in non-cardiac cells inhibits mitochondrial function and increases their vulnerability to oxidative stress without a change in mitochondrial mass.

**Figure 2.**
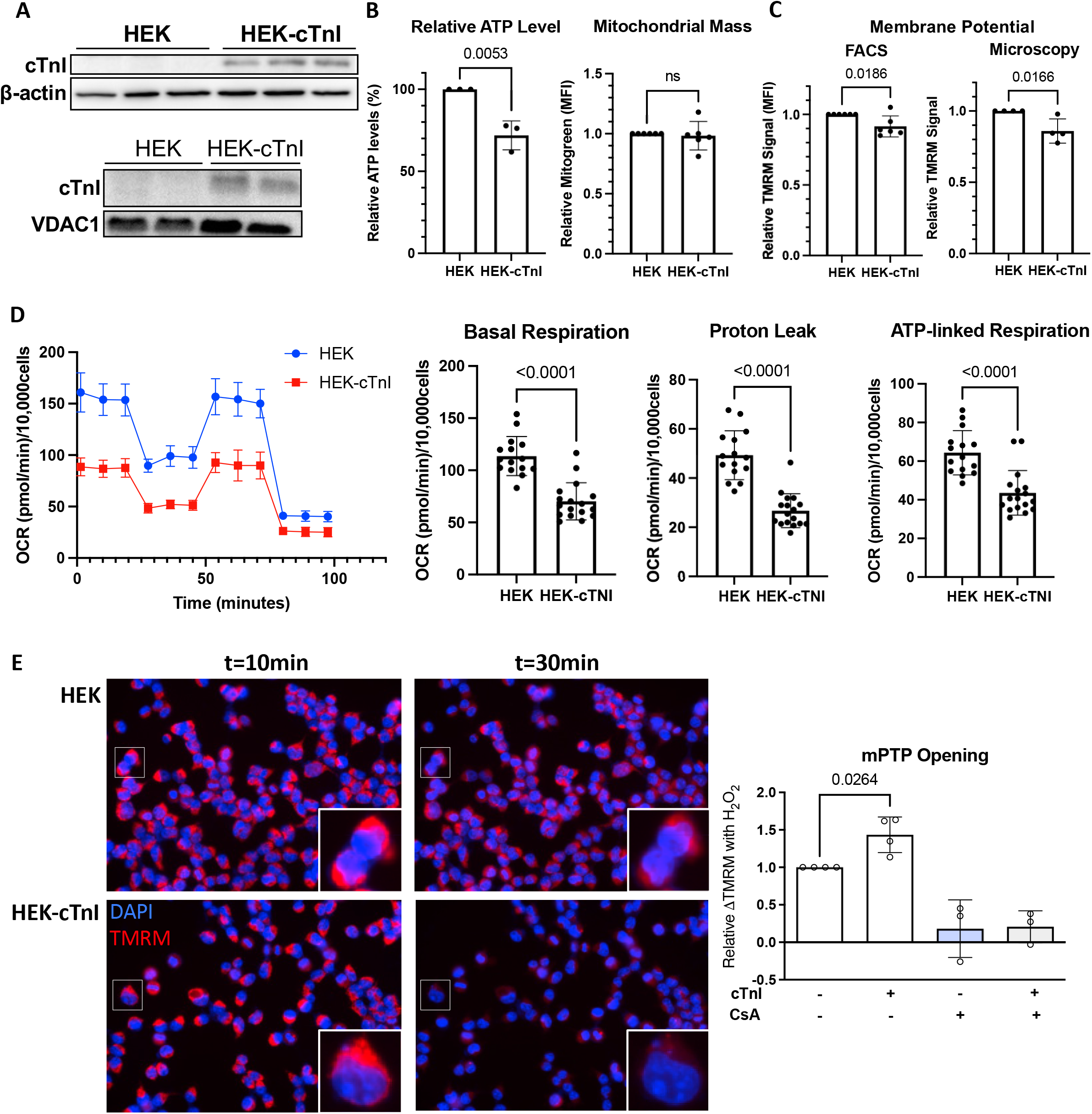
Expression of cTnI in HEK293T cells inhibits mitochondrial function. (A) HEK293T cells were engineered to stably express cTnI and compared to unmodified HEK293T cells (*top*). Mitochondria-enriched fraction shows association of cTnI with the mitochondrial fraction (*bottom*). (B) HEK cells expressing cTnI (HEK-cTnI) have decreased basal ATP levels as seen in a luciferase assay (n=3; *left*) with unchanged mitochondrial mass (*right*) as determined by flow cytometry (n=6). (C) HEK-cTnI cells have decreased mitochondrial membrane potential by TMRM fluorescence using both flow cytometry (*left*, n=6) and fluorescence microscopy (*right*, n=4). (D) HEK-cTnI cells have decreased mitochondrial oxygen consumption. Representative Seahorse experiment on left and quantifications on right for basal respiration, proton leak, and ATP-linked respiration; n=3. (E) HEK-cTnI cells have increased rate of mPTP opening as measured by rate of loss of TMRM signal after addition of 500μM H_2_O_2_. Cyclosporine A (CsA; 1μM), an inhibitor of mPTP opening, was used as control; n=3-4. *Left:* representative image showing TMRM fluorescence 10 minutes and 30 minutes after addition of H_2_O_2_. *Right:* relative change in TMRM fluorescence between 10-minute and 30-minute time points.

### cTnI directly binds the d subunit of ATP synthase

A mass spectrometry analysis of protein interactions in the mouse heart previously suggested a potential interaction between cTnI and ATP synthase^32^. We therefore examined the possibility that cTnI binds ATP synthase and directly affects its function. ATP synthase, a complex of 18 subunits, is a key component of the oxidative phosphorylation system and is proposed to be a critical component of the mPTP^33–36^. To determine whether cTnI interacts with ATP synthase, we first used an *in silico* approach that we have successfully applied before to identify potential partners of cTnI; we often find that inducible protein-protein interactions are characterized by a short sequence of similarity between the two interacting proteins^37,38^. A search for a short homology stretch between cTnI and potential binding partners identified the d subunit of ATP synthase as a potential partner; a ten amino acid sequence **<u>A</u>**_43_S**<u>RKL</u>**Q**<u>LKT</u>**L_52_ in cTnI is 70% identical and 80% homologous to the N-terminal sequence in the d subunit of ATP synthase, **<u>A</u>**_2_G**<u>RKL</u>**A**<u>LKT</u>**I_11_ (Figure 3A). This sequence in cTnI and ATP synthase subunit d is largely conserved among all mammals (Figure 3B).

**Figure 3.**
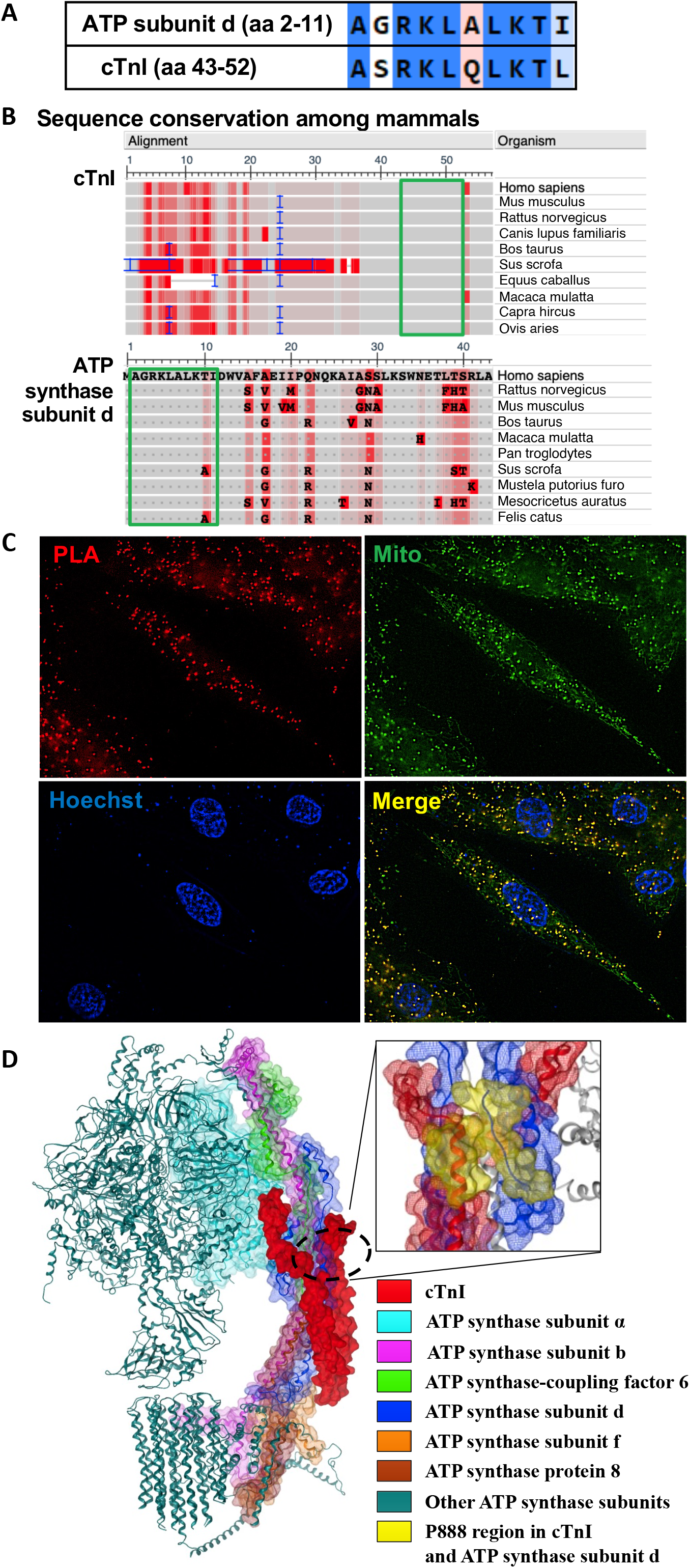
cTnI binds mitochondrial F_1_F_0_-ATP synthase. (A) Sequence homology predicted an interaction between cTnI and ATP synthase subunit d based on the presence of a 10 amino acid stretch in this otherwise unrelated protein, showing 80% similarity. (B) cTnI aa43-52 and ATP synthase subunit d aa2-11 are highly conserved in mammals. Portion of protein sequence shown. Red shading indicates frequency-based difference at each amino acid residue. Green box shows sequence of similarity between cTnI and ATP synthase subunit d. Analysis by constraint-based Multiple Alignment Tool (COBALT) plotted on NCBI Multiple Sequence Alignment Viewer. (C) A proximity ligation assay in H9c2 cardiomyocytes reveals an interaction between cTnI and ATP synthase subunit d as strong punctuate signals (red foci) were observed and overlapped with mitochondria (green). A selective rationally-designed inhibitor of this interaction was generated – P888, which mimics the cTnI-binding sequence of ATP synthase subunit d. (D) Best model for the molecular docking of cTnI in ATP synthase complex at ATP synthase subunit d site is shown. cTnI is shown as red surface and the proteins in ATP synthase complex interacting with cTnI (within 4.5Å distance) are shown as colored surfaces (ATP synthase subunit a in cyan, ATP synthase peripheral stalk-membrane subunit b in pink, ATP synthase-coupling factor 6 in green, ATP synthase subunit d in blue, ATP synthase subunit f in orange and ATP synthase protein 8 in brown). All other proteins in ATP synthase complex are shown as teal-colored ribbons. Box on the left is a magnified scope showing the relative positions of short homology stretch in cTnI (**<u>A</u>**_43_S**<u>RKL</u>**Q**<u>LKT</u>**L_52_) and ATP synthase subunit d (**<u>A</u>**_2_G**<u>RKL</u>**A**<u>LKT</u>**_11_) in the model, corresponding to peptide P888 sequence (yellow).

To determine if there is direct binding between cTnI and ATP synthase, we next used a proximity ligation assay (PLA) in H9c2 cardiac myoblasts to visualize this protein-protein interaction at a single molecule resolution. Indeed, PLA demonstrated specific interactions between cTnI and ATP synthase subunit d in mitochondria as evidenced by punctate foci overlapping a mitochondrial stain (Figure 3C). Since ATP synthase subunit d is located in the matrix sector of the peripheral stalk, the PLA provides support for occurrence of the cTnI-ATP synthase subunit d protein-protein interaction in the mitochondrial matrix. In addition, a molecular docking simulation of cTnI on ATP synthase via subunit d, using Molecular Operating Environment (MOE) software, further suggests that cTnI may bind in the peripheral stalk region of ATP synthase (Figure 3D).

### cTnI increases mitochondrial ATP hydrolysis and decreases ATP synthesis

To determine if cTnI binding to ATP synthase affects ATP synthase activity, we used an immune-captured ATP synthase-based assay. We first confirmed that recombinant cTnI bound to ATP synthase and this binding is saturable (Figure 4A). A 10-amino acid peptide corresponding to the cTnI interaction site in ATP synthase subunit d, (Figure 3A), which we termed P888, abolished cTnI binding to the immune-captured ATP synthase (Figure 4A). We then showed that cTnI binding doubled the ATPase activity (Figure 4B), and P888 blocked the cTnI-induced increase in ATPase activity (Figure 4B-C; IC_50_=0.95 μM), supporting the conclusion that cTnI binding of ATP synthase increases the activity of this complex.

**Figure 4.**
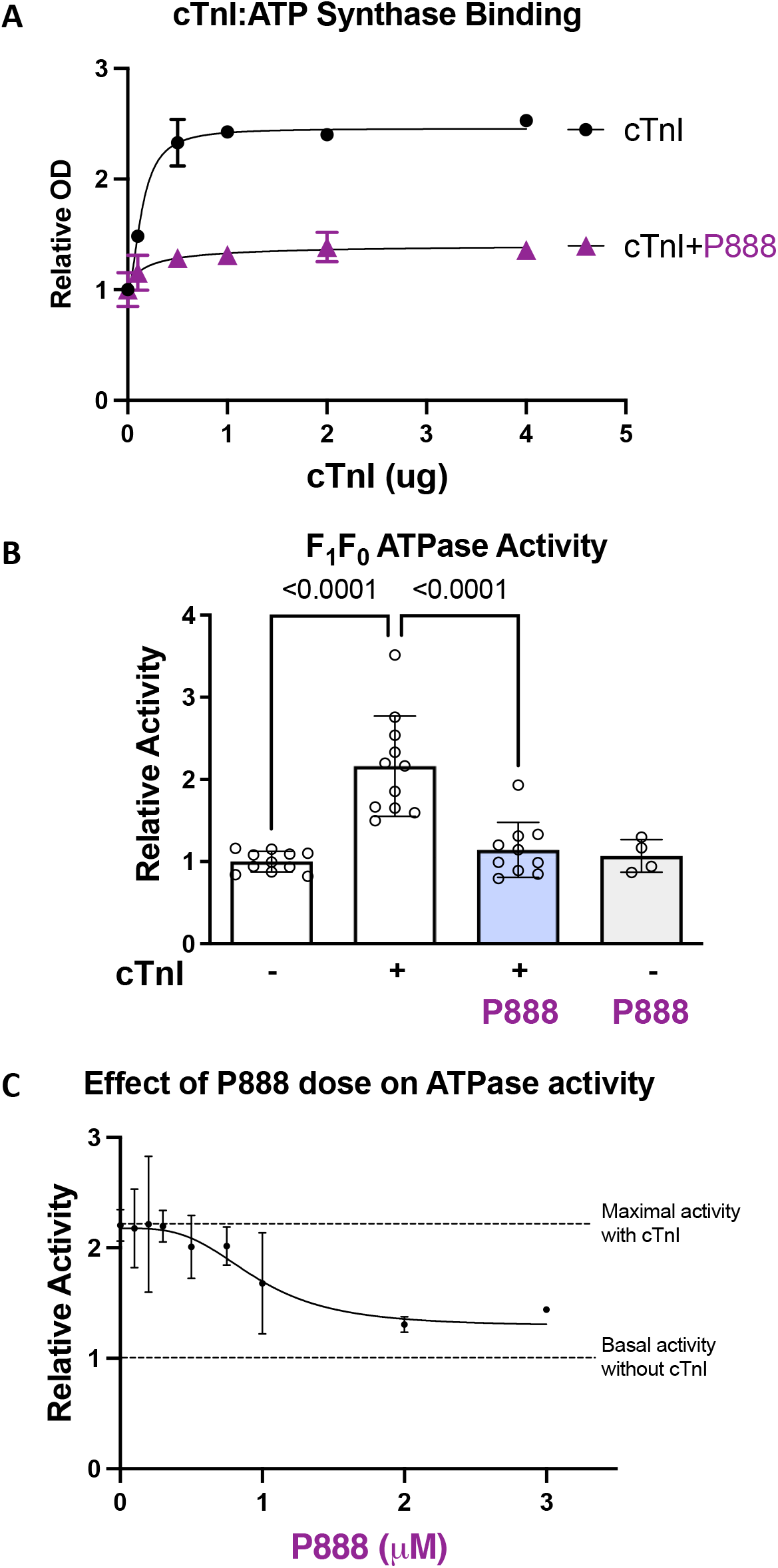
P888 modulates cTnI-ATP synthase binding and ATPase activity *in vitro*. (A) cTnI demonstrates a dose-dependent binding to ATP synthase by ELISA; binding is inhibited in the presence of P888 (2μM); n=3. (B) ATPase activity increases in the presence of cTnI, and P888 co-incubation prevents it; n=3 independent experiments conducted in 3-4 replicates each. (C) Dose-dependent inhibition by P888 of cTnI’s (2μg) effect on ATPase activity, with EC50 0.95μM: n=3 independent experiments.

N-terminal truncation of cTnI occurs at low levels in the normal heart^39^, increases in adaptation to hemodynamic stress^40^, and is associated with improved myocardial relaxation^41^. C-terminal truncation of cTnI occurs during IR^21^ and myocardial stunning^42^ and is associated with impaired contractile function^43^. We determined whether protein fragments of cTnI differentially affect ATPase activity. Indeed, we found that C-terminal truncated cTnI (cTnI1-193; Figure 5A, light blue) was sufficient to increase ATPase activity *in vitro* similar to intact protein (Figure 5A, white). Conversely, slow skeletal troponin I (ssTnI; Figure 5A, dark blue), which lacks the N-terminal 30-amino acid extension of cTnI, had no effect on ATPase activity. This suggests that the N-terminal region of cTnI is necessary in the modulation of F_1_F_0_-ATPase activity, whereas the C-terminus is not. Interestingly, in H9c2 myoblasts, GFP-tagged N-terminal cTnI did not co-localize with mitochondria. This may suggest that a segment of the N-terminus of cTnI is cleaved prior to mitochondrial entry, or that the GFP tag interferes with mitochondrial localization.

**Figure 5.**
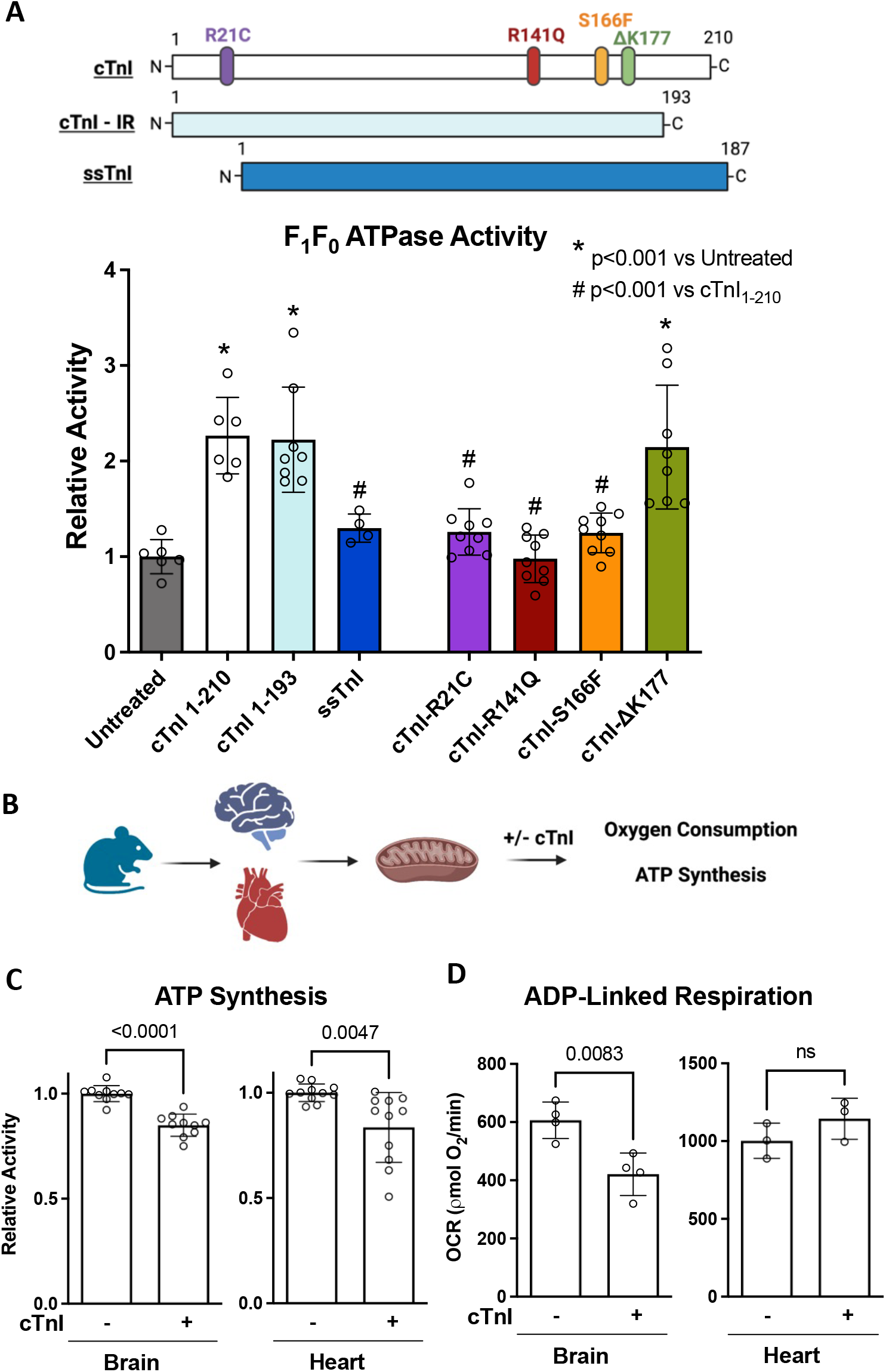
(A) Schematic of cTnI with mutations at R21C, R141Q, S166F, and ΔK177 (*top*; white), IR-induced C-terminal truncated fragment of cTnI (middle; light blue), and N-terminal truncation in ssTnI (*bottom*; dark blue). (*Left*) C-terminal truncation of cTnI (cTnI1-193) increases ATPase activity similar to full recombinant cTnI, whereas ssTnI does not. (*Right*) Mutations in cTnI differentially impact ATPase activity; n=3 independent experiments. (B) Schematic of experimental procedure: Mitochondria were isolated from mouse heart and brain and co-incubated with cTnI (4μg mitochondria:1μg cTnI), and oxygen consumption and ATP synthesis rates were measured. (C) Co-incubation of mitochondria isolated from mouse heart or brain decreases the rate of maximal ATP synthesis by luciferase assay. n=3 with 3-4 replicates each. (D) Co-incubation of isolated mitochondria with cTnI decreases ADP-linked (state III) oxygen consumption rate in brain, but not in heart; n=3-4. Figure 5B was created with BioRender.com.

Pathogenic variants in cTnI lead to cardiomyopathy with heterogeneous phenotypes (hypertrophic, restrictive and dilated) and disease course (progression rate, risk of sudden cardiac death)^44^. We determined if select pathogenic mutations in cTnI^45^ would modulate its effect on ATPase activity. Indeed, cTnI-R21C, cTnI-R141Q, and cTnI-S166F had no effect on F_1_F_0_-ATPase activity, similar to untreated ATPase complex, whereas a single codon deletion (cTnI-ΔK177) increased F_1_F_0_-ATPase activity similar to wild-type protein (Figure 5A). This suggests that pathological variants in cTnI may have differential effects on mitochondrial ATPase activity.

In respiring mitochondria, F_1_F_0_-ATP synthase catalyzes the conversion of ADP to ATP, while in hypoxic conditions where mitochondrial membrane potential is impaired, it reverses to hydrolyze ATP^46^. Given the profound effect of cTnI on ATP hydrolysis, we measured its effect on oxygen consumption and ATP synthesis in isolated mitochondria co-incubated with purified cTnI (Figure 5B). Consistent with the HEK cell data (Figure 2B, C), cTnI treatment decreased the rates of ATP synthesis (Figure 5C) in mitochondria from mouse hearts and brains, and ADP-dependent respiration in mitochondria from mouse brains, but not mouse hearts, (Figure 5D). This suggests that cTnI has distinct and opposing effects on ATP synthesis and hydrolysis by F_1_F_0_-ATP synthase complex.

### A peptide inhibitor of cTnI-ATP synthase interaction prevents IR-induced injury *in vivo*

We next reasoned that if cTnI regulates mitochondrial function *in vivo* by directly binding to ATP synthase, this effect will be inhibited by P888, the peptide inhibitor of this interaction. We therefore assessed the effect of P888 treatment on the outcome of myocardial IR injury in rats subjected to transient (30 minutes) ligation of the left anterior descending (LAD) coronary artery. At reperfusion, rats were injected intraperitoneally with TAT peptide that enables biological membrane crossing^47^ or TAT-P888 (single injection, 3mg/kg; Figure 6A). Serum lactate dehydrogenase (LDH) levels measured 3 days after reperfusion increased with IR, and this was prevented with P888 treatment (Figure 6B). Three days after IR, isoproterenol echocardiography demonstrated a decrease in systolic function as measured by fractional shortening in the vehicle-treated IR group compared to sham surgery controls; this decrease was almost completely blocked in rats treated with TAT-P888 at reperfusion only. These data suggest that cTnI’s interaction with ATP synthase at reperfusion contributes to cardiac IR injury (Figure 6C).

**Figure 6.**
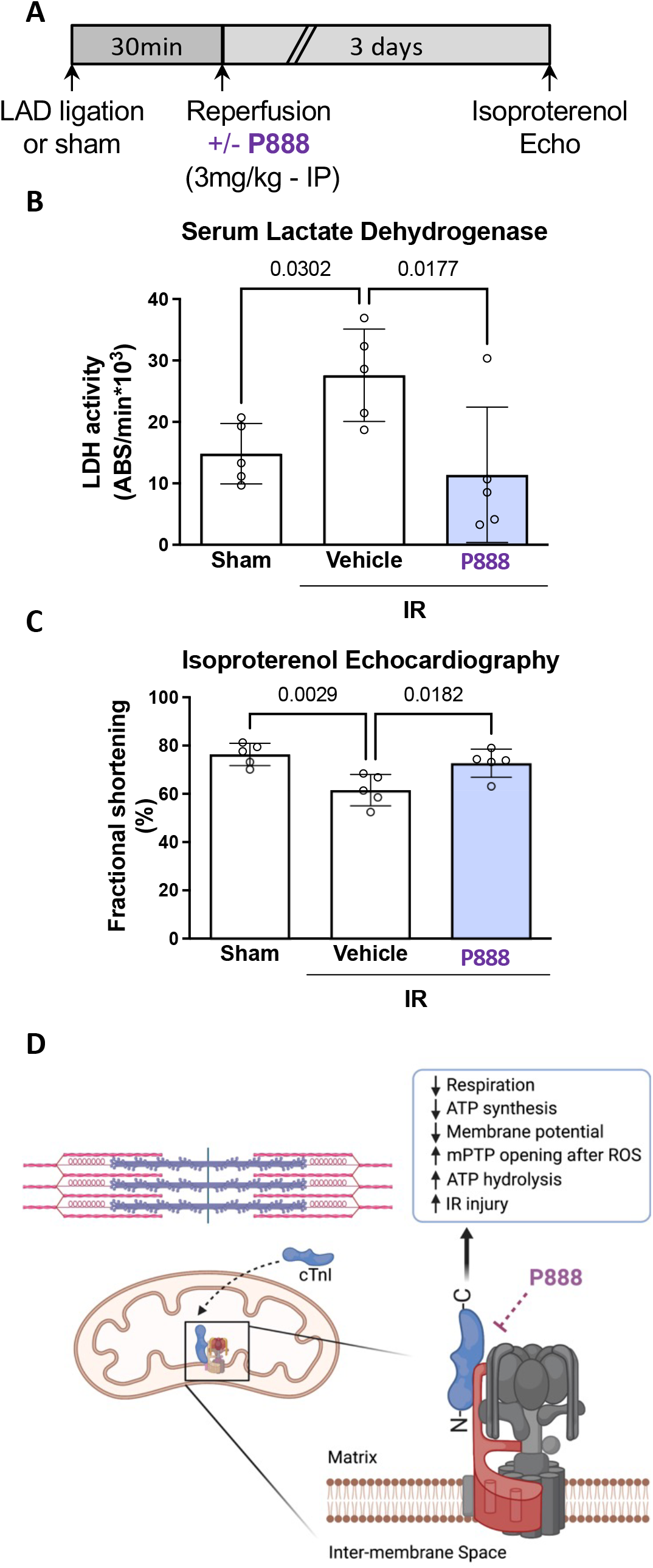
P888 treatment prevents IR-induced injury *in vivo* in rats. (A) Time course of *in vivo* myocardial IR in the presence of vehicle (TAT) or TAT-P888 treatment; n=5 animals/group. (B) TAT-P888 prevents IR-induced cardiac injury as seen by a reduction in LDH compared to vehicle. (C) Echocardiography in the presence of isoproterenol three days after IR demonstrated a decrease in systolic function, as measured by fractional shortening, in vehicle-treated animals. This reduction was prevented by treatment at time of reperfusion with TAT-P888. (D) cTnI localizes to mitochondria, binds ATP synthase, decreases mitochondrial respiration and membrane potential, increases mPTP opening after reactive oxygen species (ROS) exposure, and increases IR injury. Figure 6D was created with BioRender.com.

## Discussion

Cardiac troponin I regulates cardiomyocyte contraction in conjunction with troponin C and troponin T *via* the calcium-mediated interaction between actin and myosin. Here we identified a novel role of cTnI in mitochondria: the inhibition of mitochondrial respiration by direct binding of cTnI to the d subunit of ATP synthase. This conclusion is based on the following evidence: First, cTnI is associated with mitochondria in the heart, in cardiac myoblasts, and in non-cardiac HEK cells expressing cTnI. Second, a proximity ligation assay indicates that cTnI likely binds the ATP synthase d subunit. Third, expression of cTnI in non-cardiac HEK cells inhibited mitochondrial function as evidenced by impaired oxidative phosphorylation, a 30% decrease in ATP level, a decrease in mitochondrial membrane potential and increased sensitivity to oxidative stress induced by H_2_O_2_ treatment. Fourth, cTnI bound to immune-captured ATP synthase in a saturable manner and P888, a peptide corresponding to the sequence homology between ATP synthase subunit d and cTnI, blocked this binding at IC_50_=0.95 μM. Fifth, the functional consequences of the cTnI-ATP synthase interaction was demonstrated *in vitro*; recombinant cTnI increased the hydrolytic activity of immune-captured F_1_F_0_ ATPase, which was inhibited by P888, and treatment of isolated mitochondria with cTnI decreased mitochondrial respiration and ATP synthesis. Sixth, the N-terminus of cTnI was necessary to affect ATPase activity, and several pathogenic mutants of cTnI and the skeletal isoform of troponin I did not increase ATPase activity. Finally, myocardial IR-induced injury after transient coronary artery occlusion in rats was prevented with TAT-P888 treatment at time of reperfusion. The acute protection resulted in inhibition of the IR-induced decrease in systolic function as measured three days after infarction. Together, our work identifies a cTnI interaction with ATP synthase and the negative functional consequences of this interaction in IR-induced injury (Figure 6D).

The F_1_F_0_ ATP synthase is a protein complex with dual roles. It is a key component of oxidative phosphorylation in the production of ATP in respiring mitochondria. In addition, ATP synthase may serve as a key component of the mPTP, opening in response to cellular stress leading to cell death^33–36^. A protein-protein interaction between cTnI and ATP synthase subunit a (ATP5f1a) was previously observed in a crosslinking mouse heart proteomics dataset^32^. However, the consequence of this interaction on cardiac physiology was not assessed. Here we found that HEK cells stably expressing cTnI have decreased oxidative phosphorylation under basal conditions and increased mPTP opening following H_2_O_2_ treatment. We also found that cTnI increases mitochondrial F_1_F_0_ ATPase activity *in vitro* and decreases ATP synthesis in isolated mitochondria. In healthy respiring mitochondria, high membrane potential favors ATP synthesis by F_1_F_0_ ATP synthase, but when respiration is compromised, membrane potential decreases and ATP synthase can reverse to hydrolyze ATP^48–50^. It is possible that cTnI may function to inhibit ATP synthase and mitochondrial functions in respiring mitochondria (under basal conditions) and increases ATP hydrolysis during conditions of hypoxia. It is also possible that cTnI may affect the stability of the ATP synthase complex under stress conditions. Regardless, mitochondrial cTnI clearly exacerbates mitochondrial dysfunction and increases cell injury during excessive stress, such as during acute myocardial infarction.

Mutations along the *TNNI3* gene (which encodes cTnI) result in significant phenotypic variability, with most mutations associated with hypertrophic cardiomyopathy, while others cause dilated or restrictive phenotypes^44^. The factors leading to this phenotypic heterogeneity are not well-understood, and the avenues of targeted therapy for familial cardiomyopathies with *TNNI3* mutations are limited^51^. *TNNI3* mutations are associated with mitochondrial abnormalities in mouse models through unknown mechanisms. Here, we show that genetic variants of *TNNI3* have heterogeneous effects on F_1_F_0_-ATPase activity. The downstream effects of *TNNI3* mutation on mitochondrial function beyond ATPase activity and the role of this effect in disease pathogenesis are areas of ongoing investigation in our lab.

N-terminal truncation of cTnI occurs at low levels in normal hearts and increases in response to stress (e.g., IR injury, beta-adrenergic activation, and aging). Overexpression of this fragment (lacking cTnI aa 1-30) improves myocardial relaxation^39–41^. Interestingly, differential expression of N-terminal cTnI fragments has been shown to occur in stunned myocardium of patients after coronary artery bypass graft^52^. Similarly, in transgenic mice, expression of the slow skeletal isoform of troponin I (ssTnI, which lacks the cardiac-specific N-terminal region) in the heart improved metabolic efficiency in heart failure^53^ and IR injury^54,55^ and provided protection from endotoxemia^56^. Here, we show that unlike cTnI, ssTnI does not increase ATP hydrolysis. It is thus possible that the beneficial effect of N-terminal truncation could be at least in part due to abrogation of cTnI’s effect on mitochondrial function, supporting a damaging effect of cTnI’s N-terminal extension on mitochondria. We suggest that the novel protein-protein interaction elucidated in our study between ATP synthase and cTnI can be utilized as a potential drug target to reduce ischemic injury and sequalae of myocardial infarction and to address pathologies associated with mutations in cTnI.

## Materials and Methods

### Animals

The Administrative Panel on Laboratory Animal Care at Stanford University (protocol #10363) and the Ethic Committee on Animal Use of the Institute of Biomedical Sciences at University of Sao Paulo (protocol #60/2017) approved all animal protocols.

### Cardiac Ischemia Reperfusion – *in vivo* LAD Coronary Artery Ligation Rat Model

Myocardial infarction was induced by ligation of the left anterior descending (LAD) coronary artery for 30 minutes, as previously described^57^. Briefly, male Wistar rats were anesthetized with 3% isoflurane and intubated with a rodent ventilator at 80 breaths/minute. Body temperature was maintained at 37°C using an appropriate heating blanket. Left thoracotomy between the fourth and fifth ribs was performed, and the LAD coronary artery was ligated close to its origin from the aortic root. The normoxia control animals (sham) were exposed to the same procedure with no ligation. The free ends of the ligature were used to form a noose around a syringe plunger which was placed flat on the myocardium. Coronary occlusion was achieved by tightening the noose around the plunger for 30 minutes. Occlusion was determined by immediate blanching of the infarcted area and reperfusion was achieved by release of the ligature just after injecting an intraperitoneal injection (IP) 3mg/kg of the respective peptides. Serum lactate dehydrogenase level was measured at 3 days after MI by commercially available kit following manufacturer’s instructions (Eton Bioscience, San Diego, CA, USA). Fractional shortening was determined under basal conditions and after isoproterenol stimulation (10μg/kg, IP) at 3 days after MI by M-mode echocardiography (Acuson Sequoia model 512 echocardiographer equipped with a 14-MHz linear transducer). Treatments were performed with TAT (molecular weight: 1559.83 g/mol; CAS Number: 191936-91-1) and TAT-P888 (TAT-AGRKLALKTI; N-terminus acetylated C-terminus amidated; molecular weight: 2854 g/mol) at 1μM where denoted.

### Cell Culture and Treatments

H9c2 rat cardiac myoblasts and human embryonic kidney (HEK293T) cells were used to determine the effect of human cTnI on cellular and mitochondrial function. H9c2 cells were transfected with a cTnI construct with GFP on N-terminus and mCherry on C-terminus. A HEK293T cell line stably expressing human cTnI was generated through retroviral infection. All cells were grown in DMEM with L-glutamine, 4.5g/L glucose and sodium pyruvate (Corning) and 10% FBS.

### Cellular ATP Assay

Cellular ATP levels were determined using the ATP detection kit (Cayman Chemical, #700410) according to the manufacturer’s protocol. ATP levels detected were normalized to the total protein levels.

### cTnI:ATP Synthase Binding by ELISA

Microplates with plate-bound antibody to ATP synthase (Abcam ab109714) were co-incubated with 4μg of solubilized rat heart mitochondria (Abcam ab110347) in the presence of recombinant human cTnI tagged with GST (2μg) and P888 peptide (2μM). After washes with assay solution, plates were incubated with HRP anti-cTnI antibody (Abcam ab24460) for one hour, followed by washes, and read after addition of TMB substrate solution (Thermo Fisher N301) followed by 2N sulfuric acid. Absorbance was then quantified by spectrophotometer plate reader at 450nm.

### F_1_F_0_-ATPase Activity Assay

ATP synthase activity was measured using microplate assay (Abcam ab109714) according to manufacturer protocol. Plate-bound antibody to ATP synthase was co-incubated with 4μg of solubilized rat heart mitochondria (Abcam ab110347) in the presence or absence of recombinant human cTnI (2ug) and P888 peptide (2μM), and absorbance was measured in a spectrophotometer plate reader over a period of 120 minutes. Oligomycin treatment was used as negative control.

### Microscopy

H9c2 rat cardiac myoblasts were stained with Hoechst (1:10,000 in staining media) and MitoTracker Deep Red (1.6:10,000 in staining media) for 30 minutes in the dark. Fluorescence images were acquired using an All-in-One Fluorescence Microscope BZ-X700 (Keyence). ImageJ was used to quantify parameters from images obtained, and pseudo-color was used to better visualize co-localization.

### Mitochondrial ATP Synthesis Rate Assay

Mitochondria were isolated as described below from mouse tissue. ATP synthesis rate was determined using the luciferin/luciferase-based ATP bioluminescence Assay Kit CLS II (Roche) with modifications. Mitochondria were suspended in buffer with excess substrate (5mM pyruvate/malate). Measurements were started immediately by adding luciferin/luciferase and ADP (0.5mM final) in luminescence plate reader. The initial slope of increase in ADP-supported luciferase chemiluminescence was used to determine the rate of ATP production after subtraction of background. Oligomycin was used as control to determine the rate of non-mitochondrial ATP production.

### Mitochondrial Isolation

Male wild-type C57Bl/6 mice age 10-12 weeks old were anesthetized by isoflurane inhalation with confirmation by toe pinch reflex, followed by cervical dislocation, and excision of heart and brain. Mitochondria were isolated from heart and brain tissue using mannitol-sucrose (MS) buffer with protease inhibitor. Tissue was homogenized using a dounce homogenizer and spun at 800g for 10 minutes. Total lysate was saved at this point. The remaining supernatant was spun at 10,000g for 20 minutes to separate out the mitochondrial pellet from the cytosolic fraction. The mitochondrial pellet was washed with MS buffer and spun down at 10,000g for 5 minutes. When used for western blot, the final pellet was resuspended in MS buffer with 1% Triton-100X.

### Mitochondrial Membrane Potential and Mitochondrial Mass

Mitochondrial membrane potential and mitochondrial mass were assessed using tetramethylrhodamine, methyl ester, perchlorate (TMRM) and MitoGreen FM, respectively. HEK cells were incubated with TMRM (25 nM) or MitoGreen (100 nM) for 20 minutes in medium without serum at 37°C. Cells were washed with PBS, detached and analyzed by flow cytometry using a Cytek DxP10 flow cytometer (Cytek Biosciences, Inc.) Membrane potential was also measured by fluorescence microscopy as described below.

### Mitochondrial Permeability Transition Pore (mPTP) Opening Assay

mPTP opening was measured as previously described^58^. In brief, after staining with TMRM and Hoechst as above, live cell microscopy was completed using BZ-X700 (Keyence) in a 37°C 5% CO_2_ chamber. Relative fluorescence intensity was used to measure mitochondrial membrane potential, and after 500μM H_2_O_2_ was added to cells, the slope of decrease of TMRM signal over a 30-minute period was used to measure the relative rate of mPTP opening. Fluorescence intensity quantification was done in ImageJ.

### Molecular Docking of cTnI in ATP Synthase Complex

PDB ID: 4Y99 was loaded into Molecular Operating Environment (MOE) 2019.0102 software and 4Y99.C chain (cTnI) was kept and other chains were deleted. 4Y99.C was prepared using QuickPrep functionality in MOE at default settings. QuickPrep optimizes the H-bond network and performs energy minimization on the system. PDB ID: 6J5I [34] (ATP synthase complex) was also loaded in MOE 2019.0102 and all the ligands and metals were removed and the system was prepared using QuickPrep functionality as above. 6J5I.d chain (ATP5H) and other chains within 4.5 Å distance from 6J5I.d were kept whereas all other chains were deleted. 6J5I.d and all other chains of 6J5I within 4.5 Å distance from 6J5I.d were defined as receptor and 6J5I.d was defined as the dock site. 4Y99.C was docked in the receptor using Protein-Protein dock functionality in MOE 2019.0102 at default settings. The lowest energy docking pose of cTnI in ATP synthase complex is reported in this study.

### Peptide Synthesis

Peptide was synthesized as previously described^59^. In brief, P888 was synthesized on solid support using a fully automated microwave peptide synthesizer (Liberty, CEM Corporation) using homology sequence. For *in vivo* studies, P888 was conjugated to TAT47-57 carrier peptide, a short positively charged peptide that is used as a carrier for the delivery of the peptide into the cell, using a Gly-Gly spacer. The peptide was synthesized by SPPS (solid phase peptide synthesis) methodology with a fluorenylmethoxycarbonyl (Fmoc)/tert-butyl (tBu) protocol. The final cleavage and side chain deprotection was done manually without microwave energy. Peptide was analyzed by analytical reverse-phase high-pressure liquid chromatography (RP-HPLC) (Shimadzu, MD, USA) and matrix-assisted laser desorption/ionization (MALDI) mass spectrometry (MS) and purified by preparative RP-HPLC (Shimadzu, MD, USA).

### Protein Sequence Conservation Analysis

Mammalian protein sequences were obtained from NCBI Orthologs tool for cTnI (*TNNI3*) and ATP synthase subunit d (*ATP5H*). Alignment was conducted using NCBI Constraint-Based Multiple Alignment Tool (COBALT) (https://www.ncbi.nlm.nih.gov/tools/cobalt/re_cobalt.cgi) and visualized using the NCBI Multiple Sequence Alignment Viewer to show Frequency-Based Differences in sequences.

### Proximity Ligation Assay (PLA)

PLA was performed according to manufacturer protocols using the Sigma Duolink^®^ In Situ Red Starter Kit Mouse/Rabbit (DUO92101) and as previously described^60,61^. Images were taken at 60x or 100x using the method described in Microscopy.

### Recombinant Protein Expression and Purification

cTnI was expressed using BL 21 (DE3) *E. coli* strain transformed with pET28 plasmid containing full length human cTnI sequence or C-terminal truncated cTnI_1-193_ cloned downstream of T7 promoter in frame with N-terminal 6xHis and thrombin cleavage site. *E. coli* cells were grown in LB media supplemented with 50 μg/mL Kanamycin at 37 °C in a shaking incubator (200 rpm) until the optical density (OD_600_) of the culture reached 0.6. The culture was then induced with 0.5 mM IPTG and grown for 4 hours. Cells were harvested by centrifugation, and pellet was resuspended in lysis buffer (50 mM Tris pH 8 and 150 mM NaCl) and sonicated. Whole cell lysate was then centrifuged at 13000 rpm to separate the soluble fraction from cell debris, and clarified lysate was loaded into Ni-NTA Agarose (Qiagen, USA) gravity column pre-equilibrated with lysis buffer. The column was then washed with wash buffer (50 mM Tris pH 8, 150 mM NaCl and 40 mM Imidazole). Column bound HT-cTnI was then eluted using elution buffer (50 mM Tris pH 8, 150 mM NaCl and 400 mM Imidazole). The eluent was buffer exchanged to storage buffer (50 mM Tris pH 8 and 150 mM NaCl) using Zeba spin desalting column 7K (ThermoFisher Scientific, USA) and was flash frozen and stored at −80°C. Protein identity and purity was determined using SDS-PAGE and western blots. Protein concentration was measured using BCA assay kit (ThermoFisher Scientific, USA).

### Seahorse Assay

Cells were plated in a Seahorse XF24 Cell Culture Microplate (Agilent). Cells were washed twice with Agilent Seahorse XF Media (Agilent) supplemented with 1 mM pyruvate, 2 mM L-glutamine, and 2 mM D-glucose. Cells were then incubated in a 0% CO_2_ chamber at 37 °C for 1 hour before being placed into a Seahorse XFe24 Analyzer (Agilent). For oxygen consumption rate (OCR) and (extracellular acidification rate) ECAR experiments, cells were treated with 1 μM oligomycin, 2 μM carbonyl cyanide p-trifluoromethoxy phenylhydrazone (FCCP), and 0.5 μM rotenone/antimycin. A total of three OCR and pH measurements were taken after each compound was administered. All Seahorse experiments were repeated at least three times. For experiments with isolated mitochondria, mitochondria from mouse heart or brain (10μg) were loaded in a Seahorse XF24 Microplate with mitochondrial assay solution (MAS: 70mM sucrose, 220mM mannitol, 5mM KH_2_PO_4_, 5mM MgCl_2_, 2mM HEPES, 1mM EGTA, 0.2%BSA fatty acid-free, pH 7.4) plus pyruvate (5mM) and malate (5mM). Mitochondria were sequentially treated with substrate+ADP (2.5mM), oligomycin (2μM), FCCP (4μM), and Antimycin A (4μM).

### Western Blot Analysis

A BCA assay (Thermo Scientific, #23225) was used for protein quantification. Samples were boiled with Laemmli buffer containing 2-mercaptoethanol, loaded on SDS–PAGE, and transferred on to PVDF membrane, 0.45 μm (Bio-Rad). Membranes were probed with the antibody of interest and then visualized by ECL (0.225 mM p-coumaric acid; Sigma), 1.25mM 3-aminophthalhydrazide (Luminol; Fluka) in 1 M Tris pH 8.5. Relative densitometry was measured in FIJI ImageJ. The antibodies used in this study are: Anti-VDAC1 Antibody [20B12AF2] (Abcam; ab14734; 1:1000; lot #: GR3296736-19); β-Actin (8H10D10) Mouse mAb Antibody (Cell Signaling Technologies; #3700; 1:1000; lot #: 17); Cardiac Troponin I (Abcam; ab47003; 1:500; lot #: GR3248433-1); Cardiac Troponin C (Abcam; ab137130; 1:500; lot #: GR106421-9); HRP anti-cTnI mouse mAb (Abcam; ab24460; 1:500; lot #:GR34365881); Mouse IgG HRP linked whole Ab (Sigma; #NA931-1ML, 1:5000; lot #: 17041904), and Rabbit IgG HRP linked whole Ab (Sigma; #NA934-1ML, 1:5000; lot #: 17065614).

### Statistical Analysis

Values are presented as mean ± S.D. relative to the average value for the control group unless otherwise stated. Group differences were assessed by Student’s t-test (in experiments with two groups) or ANOVA (in experiments with more than 2 groups) with correction for multiple comparisons and with statistical significance established at *p* < 0.05 (Graph Pad Prism).

## Acknowledgments

We appreciate the informative discussions and generosity of the members of the Mochly-Rosen laboratory. We thank Dr. Fabio De Lisa for his encouragement and Dr. Paolo Bernardi for much advice and critical review of our work. Preliminary data on ATPase activity by Dr Valentina Giorgio are also acknowledged. We thank Dr Masataka Kawana and Dr James Spudich for generously providing recombinant cTnI mutant proteins. This research was supported by NIH T32 (HL09427411A1) to AE, AHA Postdoctoral Fellowship (19POST34380299) to AJL, NICHD K99 HD099387 and Maternal Child Research Institute MCHRI pilot grant to BH, FAPESP (2021/09484-7) to JCBF, and NIH grant (R01-HL052141) to DM-R.

## Author Contributions

Study conception and design were performed by AJL, AE and DM-R. The animal/cell experiments, sample collection, and subsequent experimental analysis were conducted by AE, AJL, VV, SP, LRGB, NPO, BBQ, JCC, and JCBF. The data analyses were performed by AE, AJL, VV, SP, LRGB, NPO, BBQ, JCC, and JCBF. The manuscript was drafted by AE, AJL, and DM-R. All authors read and approved the final manuscript.

